# Design of MEG-Based Brain-Machine Interface Control Methodology Through Time-Varying Cortical Neural Connectivity & Extreme Learning Machine

**DOI:** 10.1101/2022.12.05.519149

**Authors:** Caglar Uyulan

## Abstract

Human-machine interfaces contribute to the improvement of the life quality of physically disabled users. In this study, a non-invasive brain-machine interface (BMI) design methodology was proposed to control a robot arm through magnetoencephalography (MEG) based on directionally modulated MEG activity that was acquired during the user’s imagined wrist movements in four various directions. The partial directed coherence (PDC) measure derived from functional connectivity between cortical brain regions was utilized in the feature extraction process. The time-varying parameters were estimated based on a time-varying multivariate adaptive autoregressive (AAR) model, that can detect task-dependent features and non-symmetric channel relevance for mental task discrimination. An extreme learning machine (ELM), that utilizes Moore-Penrose (MP) generalized inverse to set its weights and does not necessitate a gradient-based backpropagation algorithm was employed to generate a model with the extracted feature set. The output of the task classification model was embedded into the robotic arm model for realizing control-based tasks. The classification results dictate that the proposed BMI methodology is a feasible solution for rehabilitation or assistance systems that are devised to help motor-impaired people. The proposed methodology provides very satisfactory classification performance at a fast learning speed.

## 1. Introduction

BMIs are built as communication devices, that encode brain activity to a machine command signal, not involving muscles. The utilization of BMIs is quite helpful for users having diseases or traumatic injuries which cause muscle control degradation or motor disabilities (termed as a locked-in syndrome). The locked-in syndromes may comprise a wide spectrum of diseases and traumata, i.e. amyotrophic lateral sclerosis, cerebral palsy, muscular dystrophy, multiple sclerosis, brainstem stroke, and brain or spinal cord injury.

Practical applications of brain-computer interfaces (BCIs) cover electrophysiological signals, such as electroencephalography (EEG), electrocorticography (ECoG), and MEG. BCIs generally control a robotic end system via these aforementioned signals recorded from the synchronized activity of neuron groups inside the brain of subjects. The subjects can be people having different ages, gender, and physical characteristics. The aim here is to establish a user-independent, generic model of the BMI and to search for robust-adaptive algorithms that minimize the variance of classification errors concerning the various kind of users. The combination of “feature extraction/dimension reduction/feature selection/classification” algorithms with the best classification performance and the least possible error variance concerning users becomes the main model. As a result of this main model obtained, i.e., the robot trajectory control can be realized. The strong hypothesis (model) that is generated, is expected to guarantee the mental task classification of random users by limiting their errors to a lower band. As a result of this research, the BMI can be brought to a level that competes with commercial applications. The design procedure is divided into three basic modules: data acquisition, signal processing, and machine control. The majority of brain waves along the scalp are collected, and the wave information is transferred to a central processing unit (CPU) for processing in the data acquisition process. The CPU filters the signals and runs algorithms that extract features to control the machine. MEG can decrease training time while increasing the robustness of a BMI. As compared to the EEG, MEG has a larger number of sensors. Therefore, the spatial resolution is more. The detection of the frequency information can be above 40 Hz, which is not capable in EEG recordings [1, 2].

BMI architecture to be designed: 1) should adapt to non-stationary brain dynamics, 2) should generate neural signals from general brain states independently of the users, 3) should have a generalizable structure that will allow an easy transition to control applications, and 4) should demonstrate high classification performance [3, 4, 5, 6].

EEG-based BCIs suffer the problem that the recorded signal is non-stationary, that is, the data is subject to covariate shift. The non-stationarity can occur from the transition of the training model without feedback to online use with feedback, the shift of sensors to different brain regions due to head movements in MEG, or variations in mental status over time. These non-stationary cases are valid for a classifier trained on training data showing this mismatch [7, 8].

Since the MEG is a non-stationary signal, many feature extraction methods that originated from time-domain, frequency-domain, and time-frequency domain analysis, are utilized in the literature [9, 10, 11, 12]. Time-domain features are extracted directly from the signal and focus on the (averaged) time course. Frequency-domain features highlight the signal power in frequency band components. Time-frequency domain features analyze how the power spectrum changes w.r.t. time. The short-time Fourier transform and the wavelet transform are the most applicable [13, 14, 15, 16].

Spatial filtering, which extracts signals from multiple sensors to look at the activity localized in a particular brain region, is also applied. Some of the common spatial filtering methods can be listed as; *bipolar montage*, where bipolar channels are evaluated by subtracting the signals from two collocated electrodes [17]; *common average reference*, which subtracts the average value of the full electrode montage from that of the specific channel [18]; *laplacian method*, which evaluates for each electrode location the second derivative of the instantaneous spatial voltage distribution by combining the value at that location with the values of a set of surrounding electrodes [19]; *common spatial patterns*, which analyzes multi-channel data based on savings from two tasks. The filters are utilized to optimize the variance for one task and minimize it for the other tasks [20]. Other techniques have also been utilized in electrophysiological feature extraction, such as the Kalman filter, fractal dimensions, and entropy [21, 22, 23]. The AAR algorithm based on the Kalman filtering approach has also been applied to analyze the electrophysiological signals. The parametric model has an adaptive closed-loop controller with a recursive Bayesian estimator and a linear-square regulator. It estimates states and system parameters from different mental tasks and feeds them back to the optimal controller. The performance of BMIs is enhanced by closed-loop solver adaptation or multiplicative recurrent artificial neural network (ANN) solver coupled with KF. Thanks to this solver, the learned training datasets become more robust and provide a wide variety of neural-kinematic mapping learning [24, 25, 26].

Coming to the classification stage, the use of ANNs is quite common. However, the learning rates of feed-forward ANNs, including deep learning applications, are quite low. Among the main reasons for this are the use of slow gradient-based learning algorithms in the training of ANNs and the iterative updating of all parameters of the networks in this way at each stage. ELMs have succeeded in solving this problem by proving that hidden nodes do not necessitate learning and are recursively set. ELM consists of generalized single or several hidden layer feedforward networks. ELMs give importance to feature representations in hidden layers compared to support vector machines (SVM). These models can demonstrate good generalization performance and learn extremely faster than backpropagation networks. Random input layer weights add to the generalization capabilities of a linear output layer solution as it has nearly orthogonal (weakly correlated) hidden layer properties. If the weight range is constrained, the orthogonal inputs yield a larger solution space volume with these constrained weights. Thanks to the small-weight norms, the system is more stable and robust to noise. The random hidden layer generates weakly correlated hidden layer features, proposing a solution with a low norm and strong generalization performance. Various types of ELM have been proposed in the literature [27, 28, 29].

To learn effectively from various data types, ELMs need to be modified to suit the problem. When new samples are added to the dataset and grown, the ELM is retrained, but retraining the network is inefficient because the proportion of new incoming data is small. Therefore, [30] proposed an online sequential ELM (OS-ELM). The basic idea behind OS-ELM is to avoid retraining on previous examples using a sequential method. OS-ELM reconstructs settings using new instances sequentially after startup. [31] developed an incremental ELM (I-ELM). When a new hidden node is introduced, I-ELM randomly adds nodes to the hidden layer one by one, freezing the output weights of the existing hidden nodes. I-ELM is effective for single-layer feed-forward networks (SLFN) with piecewise continuous activation functions. [32] proposed a pruned ELM (P-ELM) algorithm as a systematic and automated strategy for building ELM networks. Using a small number of hidden nodes can cause under/overfitting problems in model classification. Compared to traditional ELM, simulation results showed that P-ELM results in compact network classifiers that produce fast responses and strong prediction accuracy on unseen data. [33] proposed an error minimization-based method (EM-ELM) that determines the number of hidden nodes in generalized SLFNs by growing hidden nodes one by one or group by group. As the networks grow, the output weights are gradually changed, significantly reducing the computational burden. The simulation results in sigmoid-type hidden nodes showed that this method can greatly decrease the computational cost of ELM. Nodes are randomly added to the network until they reach a certain error value of ϵ. A new learning algorithm called an evolutionary ELM (E-ELM) has been developed to optimize input weights and latent biases and determine output weights. Output weights are determined analytically using the MP generalized inverse [34]. The stability and generalization performance of the ELM was investigated. This system consists of two stages. In the first step, a forward iterative algorithm is applied to select hidden nodes from randomly generated candidates at each step, then hidden nodes are added until the stopping criterion is matched. In the second stage, unimportant nodes are removed [35, 36]. A kernel-based ELM (KELM) inspired by SVM has been developed and the main kernel function used in ELMs is the radial basis function. KELMs are used to design Deep ELMs (DELMs) [37]. DELMs utilize KELM as an output layer [38]. Voting-based ELM (V-ELM) has been proposed to improve performance on classification tasks. In V-ELM, not just one network is trained, but many are trained and then the fittest is chosen according to the majority voting method [39].

In this study, a signal processing methodology was developed for the subjects who control the movement of a robotic arm using four motor imagery signals related to the following wrist movement states, respectively: right, forward, left, and backward. Section 2 gives a background on the MEG acquisition system and the Robot arm, followed by a discussion on the classification & control strategy implemented in this study. In Section 3, the analysis results were presented. Section 4 includes the discussion and conclusive summary part.

## 2. Materials and Methods

### 2.1. MEG data acquisition & preprocessing

The data set was provided by the Brain Machine Interfacing Initiative, Albert-Ludwigs-University Freiburg, the Bernstein Center for Computational Neuroscience Freiburg, and the Institute of Medical Psychology and Behavioral Neurobiology, the University of Tübingen collected for the BCI Competition IV [https://www.bbci.de/competition/iv/]. The data set comprises directionally modulated low-frequency MEG activity that was acquired from two healthy, right-handed subjects. They performed wrist movements in four various directions, i.e. we have four different classes. The task was to move a joystick from a center position toward one of four targets located radially at 90^°^ intervals (four-class center-out paradigm) utilizing exclusively the right hand and wrist.

The MEG device has 10 channels and 625 Hz. sampling rate. The data were band-pass filtered (0.5 to 100 Hz., Butterworth, 3^rd^ order) and resampled at 400 Hz. The data comprise signals from ten MEG channels which were positioned above the motor areas given in Fig.1.

**Figure 1.**
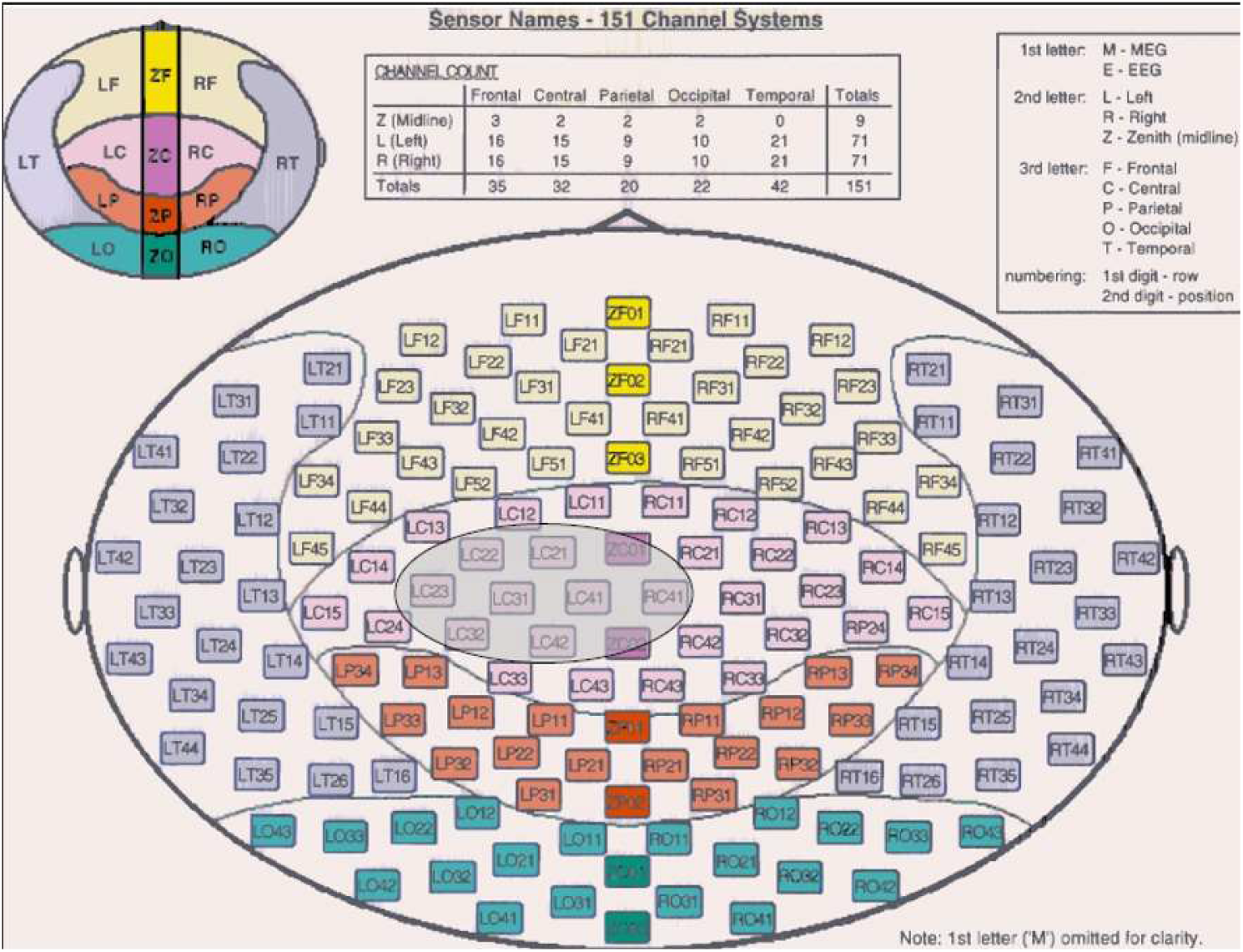
The MEG channel positions. ‘LC21’, ‘LC22’, ‘LC23’, ‘LC31’, ‘LC32’, ‘LC41’, ‘LC42’, ‘RC41’, ‘ZC01’, ‘ZC02’ [https://www.bbci.de/competition/iv/].

The details about the data acquisition and signal processing processes can be accessed at [https://www.bbci.de/competition/iv/]. Data were collected from two separate subjects including training sessions during 40 seconds/trials for each mental task, and testing sessions during 73 trials by executing random tasks. The test data will be used as external validation for assessing the performance of the proposed classifier.

The training data were merged as the size of the matrix for each mental task-based class is 40 (# *of trial*) × 800 (# *of sampling data*) × 10(# of channels). The train data matrix was reshaped into a two-dimensional form as 32000 (# *obtained from multiplication of trials with the sampling data*) × 10(# of channels). The total size of the training data is 128000 × 10.

The testing data including the random sequence of mental tasks were merged as the size of the matrix is 73 (# *of trial*) × 800 (# *of sampling data*) × 10(# of channels). The test data matrix was reshaped into a two-dimensional form as 58400 (# *obtained from multiplication of trials with the sampling data*) × 10(# of channels).

### 2.2. Feature extraction

The feature extraction process includes the transformation of the preprocessed signal into a feature matrix by attenuating noise and focusing on important data. The *AR*_*FIT*_-package is utilized to find an optimum model order for the time-varying Multi-variate autoregressive (MVAR) model. The time-invariant parameter and the order of the model can be estimated using *AR*_*FIT*_-package. “Schwarz’s Bayesian Criterion” is applied for the order estimation and then the model order is fixed for further analysis. Time-varying PDCs are evaluated through the time-varying MVAR model matched with the signal using an AAR algorithm, which uses linear Kalman filtering for parameter estimation. A surrogate data method having 50 realizations is performed to choose the most essential quantities of the measure at a 99% confidence level. Surrogates are constructed by randomizing all samples of the signal to remove the causal relations among them [40, 41]. The algorithm embeds the linear Kalman filtering to update the MVAR parameters for each time sample.

A *d*-dimensional time-varying MVAR process is given in Eq.1.

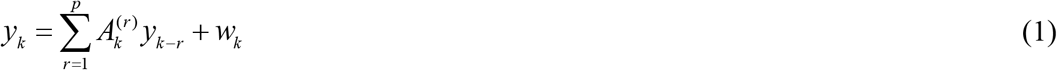

where *p* is the model order, *w*_*k*_ ∈ *R*^*d*^ is a zero-mean white process noise vector, and 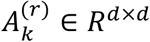is the matrix of autoregressive coefficients at each lag *r* and time point *k* = *p* + 1, …, *N*.

A form of state space equations is built from the MVAR equations by re-organizing all matrix parameters into a state vector of the dynamical system and focusing the non-stationary signal as the measurement is given in Eq.2 [42, 43].

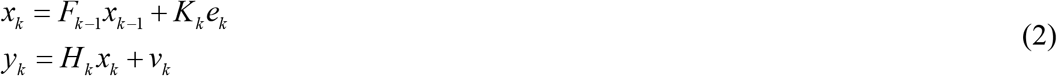

where *x*_*k*_ is the parameters vector (state vector), *K*_*k*_ is the Kalman gain, *H*_*k*_ is the measurement matrix (observation matrix), *e*_*k*_ is the one-step prediction error and *y*_*k*_ is the estimated vector.

*x*_*k*_ and *H*_*k*_ are stated in Eq.3.

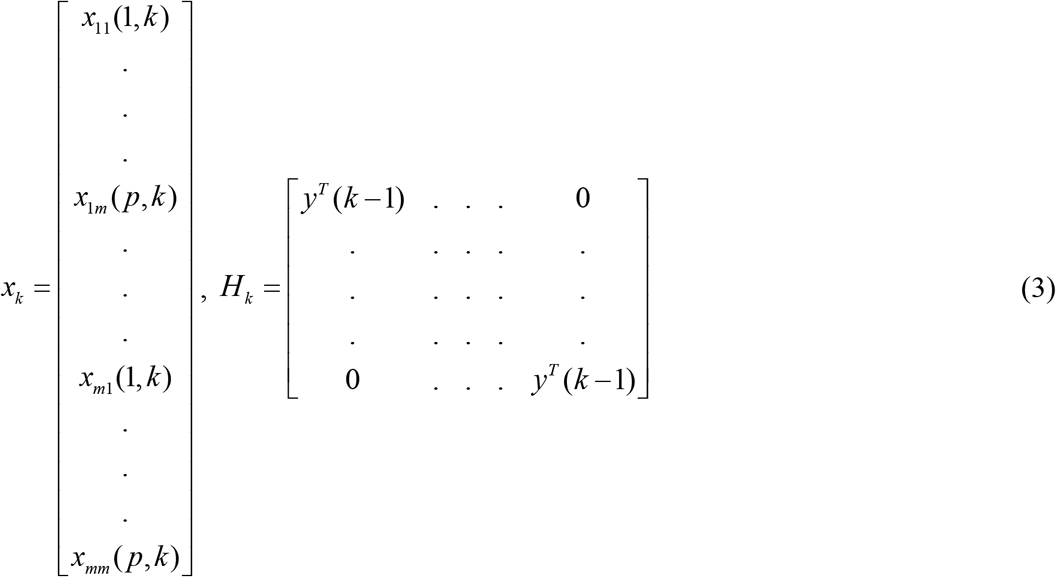

where *y*^*T*^(*k* − 1) = [*y*^*T*^(*k* − 1) … *y*^*T*^(*k* − *p*)]. The elements of the state vector are estimated via the Kalman filtering approach. The process and observation noise covariance matrices (*w*(*k*), *v*(*k*)) are updated using a specific method given in Eq.4 [44].

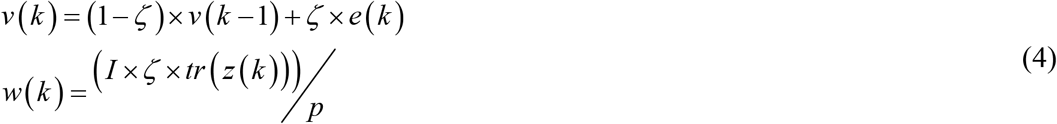

where *ζ* is the update coefficient, *z*(*k*) is the a-posteriori correlation matrix. *I* is the identity matrix, and × implies the matrix product operator.

The update coefficient determines the time resolution and the smoothing of the AR estimates. The 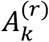 matrices given in Eq.1 are in the following form given in Eq.5.

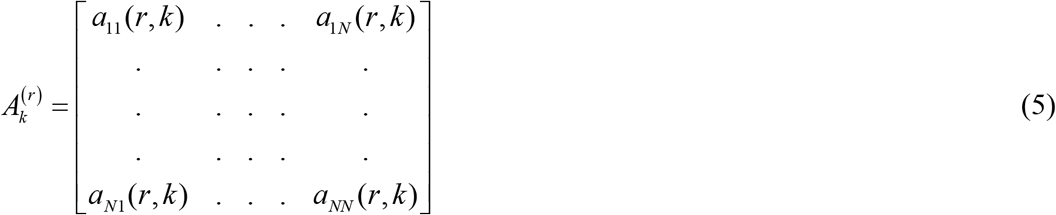

*for r* = 1, …, *p* and their elements are predicted utilizing the adaptive method given in Eq.1-4. Based on that information, adaptive time-varying connectivity measures are defined on the following Z-transform of the MVAR parameters 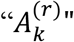 to the frequency domain as in Eq.6.

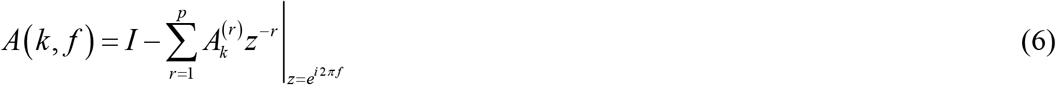

PDC, which is a time-varying connectivity measure is defined in Eq.7 [45, 46].

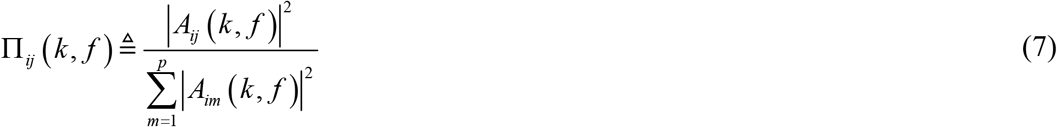

PDC quantifies the direct influence from time-series *j* to time-series *i*, after discounting the effect of all the other time series. The square exponents enhance the accuracy and stability of the estimates while the denominator part permits the normalization of outgoing connections by the inflows [47].

### 2.3. Classification

Extracted features are then defined as input neurons to the classification algorithm. The output layer should include four neurons for the four classes that represent the four wrist movement states. The number of neurons in the input layer changes according to the length of the feature vector. NNs and SVMs play key roles in machine learning and data analysis. However, it is known that there exist some challenging issues with them such as intensive human intervention, slow learning speed, and poor learning scalability. In this paper, we use ELM for the classification. ELMs are a kind of feedforward neural network, which does not necessitate gradient-based backpropagation for learning. It utilizes MP generalized inverse to set its weights. ELM not only learns up to tens of thousands faster than NNs and SVMs but also provides unified implementation for regression, binary and multi-class applications. ELM is efficient for time series, online sequential, and incremental applications. ELM is also efficient for large datasets.

A single hidden layer feedforward NN is shown in Fig.2.

**Figure 2:**
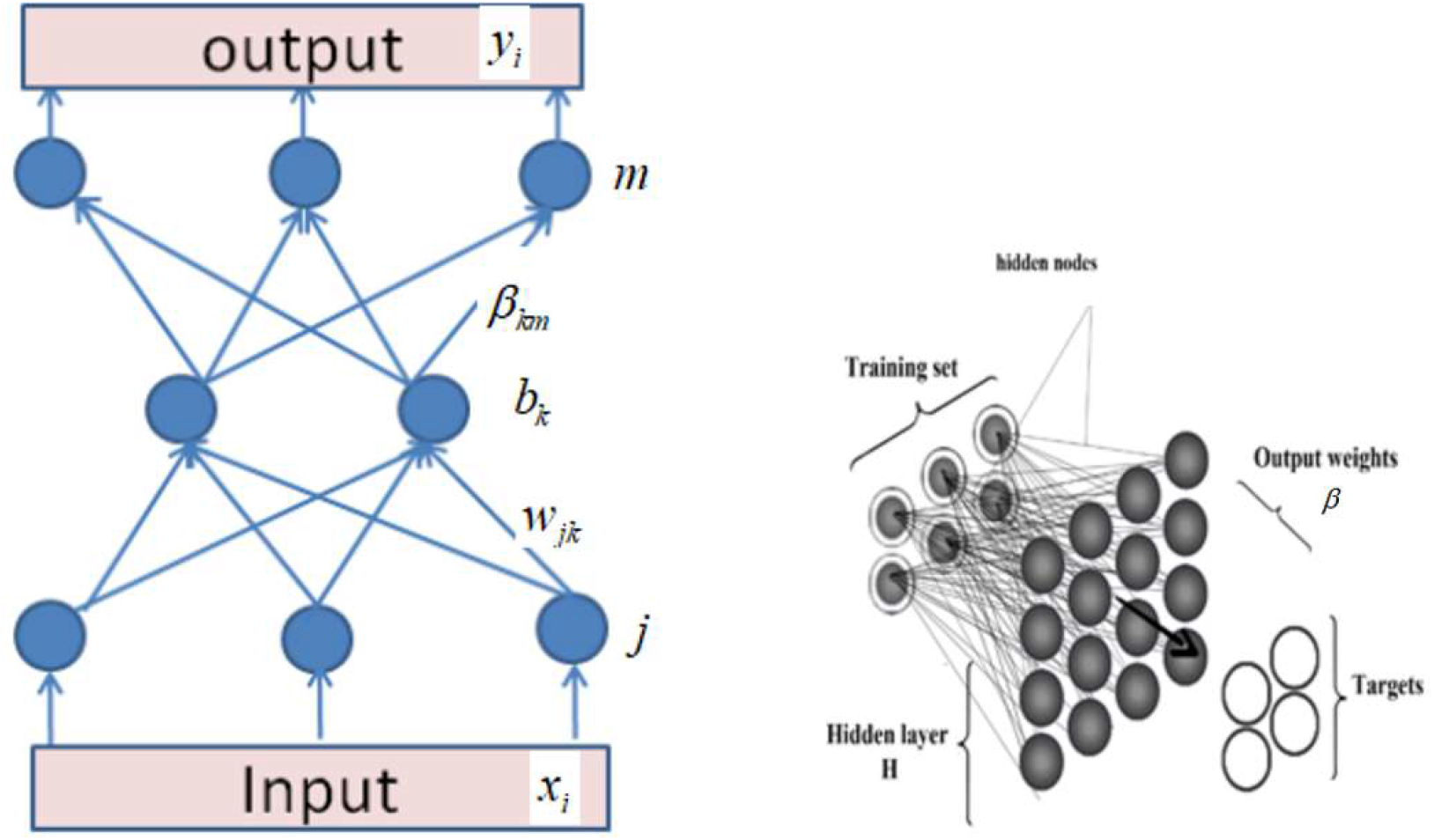
A generic single hidden layer feedforward neural network architecture [48].

The algorithm of a single hidden layer feedforward NN can be listed as multiplication inputs by weights, adding bias, implementing the activation function, repeating the first three steps with a number of layers times, evaluating output, backpropagating, and repeating every step, respectively. The ELM differs from the feedforward NN by removing the repeating steps and replacing the backpropagation step with a matrix inverse operation.

The output of the ELM is evaluated as in Eq.8.

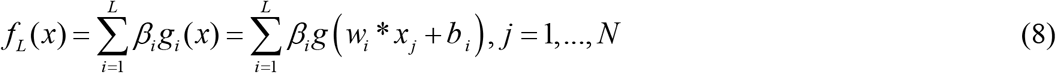

where *L* is the number of hidden units, *N* is the number of training samples, *β*_*i*_ is the weight vector between *ith* hidden layer and output, *w* is the weight vector between the input and hidden layer, *g*(.) is an activation function, *b* is a bias vector, and *x* is an input vector.

*β* is a special matrix due to the pseudo-inverse operation. Eq.8 can be represented in a compact form as in Eq.9.

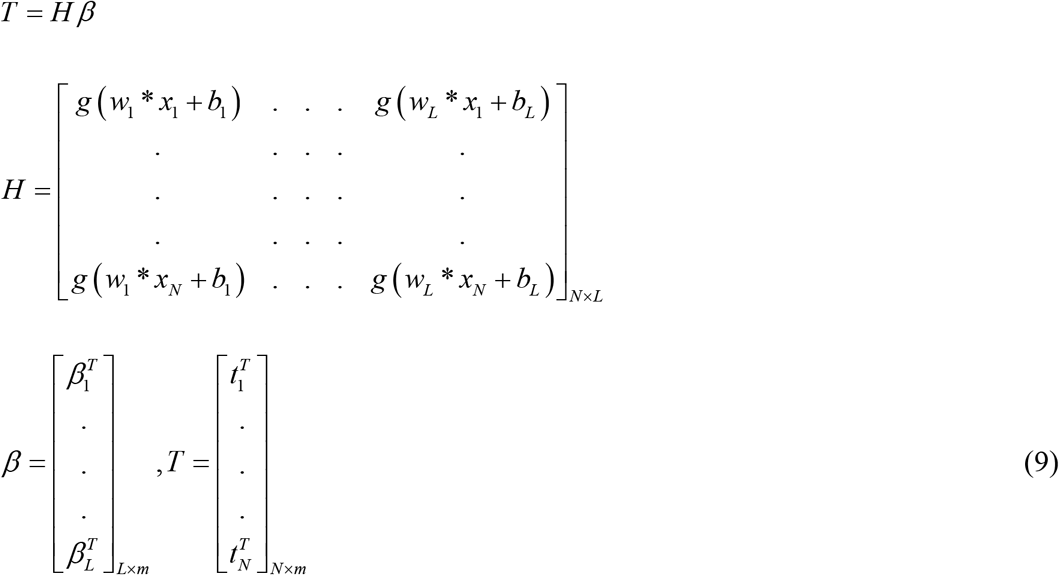

where m is the number of outputs, H is called the hidden layer output matrix, and T is a training data target matrix.

Since there are no certain rules for choosing the number of hidden neurons, the EM-ELM method was used for finding the optimal configuration.

### 2.4. Machine Interface Design

JACO robotic arm, which is a generic 6-axis robotic manipulator with a three-fingered hand can be used for the physical realization and machine interface of the proposed algorithm. The arm has six degrees of freedom in total with a maximum reach of 90 cm radius sphere and a maximum speed of 30 cm/s. It is made of three sensors: force, position, and acceleration. This arm should be suitable for a person with a disability of the upper arm and can be placed in a wheelchair. The upper arm of the robot is made of three links which are similar to the upper limb of the human body, as shown in Fig.3.

**Figure 3:**
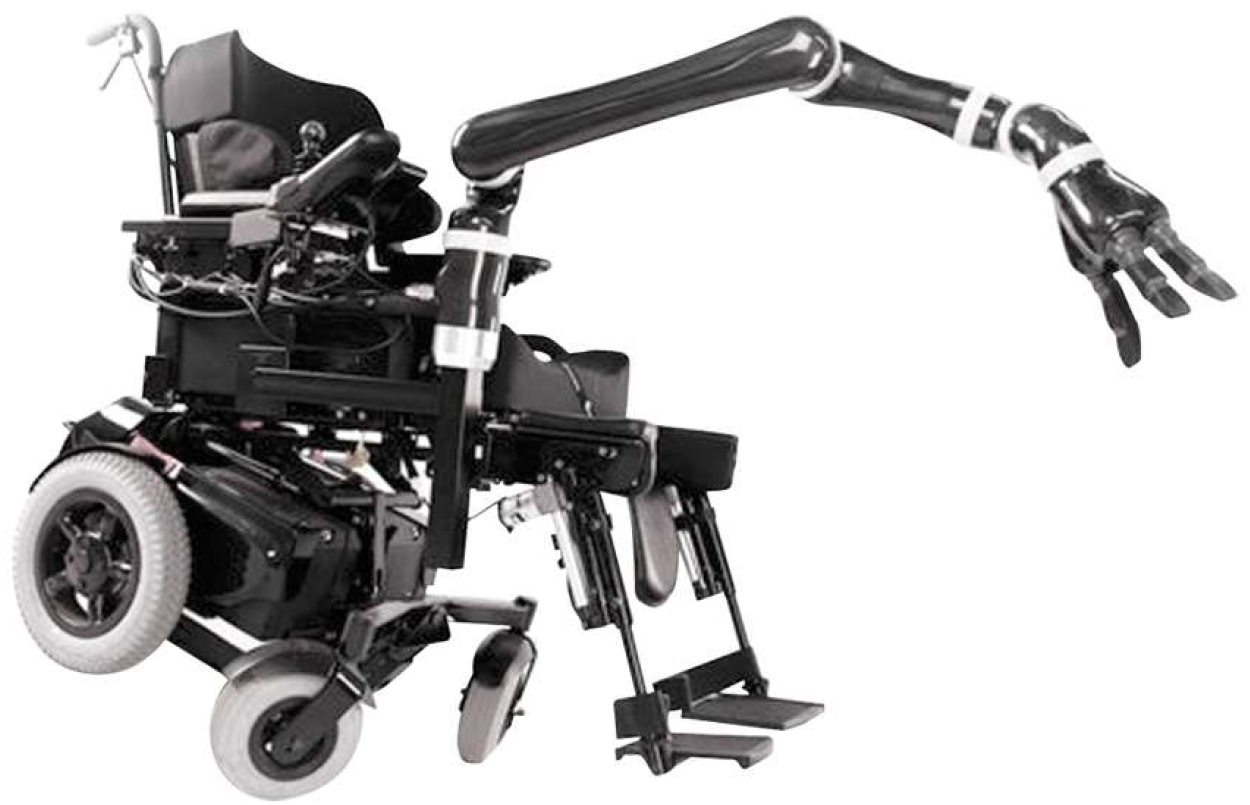
JACO robotic arm mounted on an electric wheelchair [49].

An API, which gives freedom of control to users, is provided by the manufacturers. The subject needs to control the movement of the robot arm toward a given target by using four mental commands: Forward (F), Backward (B), Left (L), Right (R), and No Movement command. To end the movement of the robot arm, the subject would generate a “No Movement” command by taking no action. The arm could move on 2 axes (x and y) and in four directions (forward (+y), backward(-y), left(+x), and right(-x)). The control signals are generated according to the mental commands. To move the arm forward the subject imagines the forward wrist movement, and performs backward wrist movement to move the arm backward. The subject imagines moving his/her right wrist to move the robotic arm right and imagines moving his/her left wrist to move the robotic arm left. Forward differential kinematics gives the relation between joint velocities and tip velocity. This relation is in matrix form called Jacobian. To form Jacobian, first propagation matrices are created. The propagation matrix given in Eq.10 produces the relation between sequenced joints.

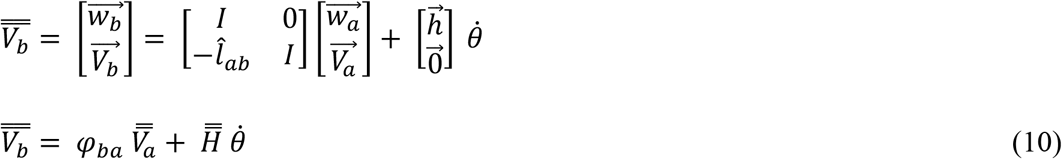

For fixed base manipulators, the robot propagation matrix can be constructed by using these joint relations. This matrix is called *φ* and gives all joints velocity vectors. To reach tip velocity, Eq.11 should be evaluated.

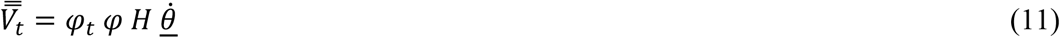

By using this equation, the robot tip velocity can be calculated concerning joint velocities. So the Jacobian is given in Eq.12.

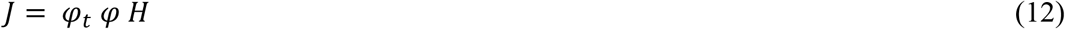

MATLAB Simulink Code is created by using the above equations. First, the jacobian is implemented, after that according to Rodriguez’s formula, the rotation matrices for each joint are built. The coordinate frames for each joint change when the robot moves. For this reason, coordinate frames are updated at each sample time by using joint velocities, sample period, and rotation axes. However, this update mechanism is correct if the joint velocity is constant along the sample period. The selected sample time is 1 millisecond. The Simulink block diagram of the robot jacobian is shown in Fig.4.

**Figure 4:**
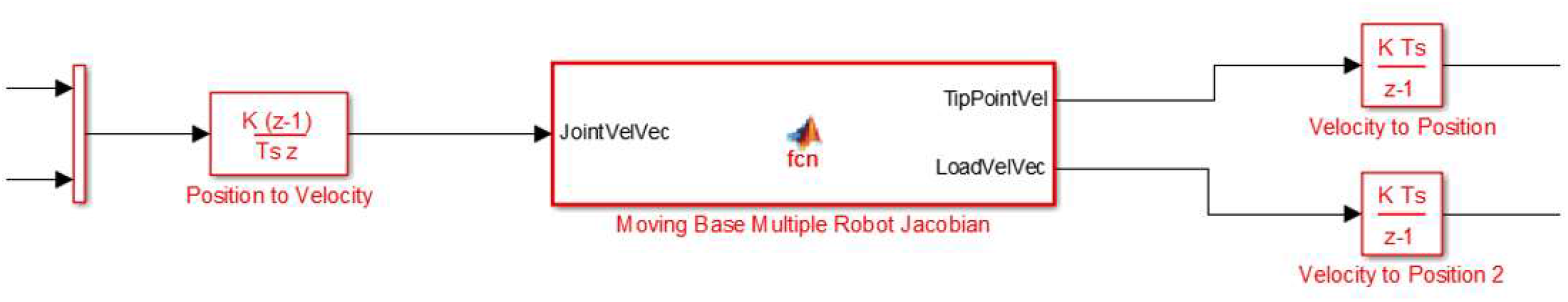
Simulink block diagram of the robot jacobian.

Inputs are Base Positions (6×1), Robot Joint Angular Positions (6×1), and Outputs are Tip Points Position, and Load Center Position. Tip and load center positions are fed to these shapes at the VRML file to confirm the operation of the moving base kinematic code. Only the angular part of the base position is used. These angles are Euler XYZ angles and converted to vector-angle representation then fed to VRML file. Two common ways to create a dynamical model of a robot are Newton-Euler-based and Euler-Lagrange-based dynamical modeling. The Newton-Euler method, which is a vectorial approach, was explained in this paper for dynamical modeling.

The dynamic model stated in Eq.13 describes the relationship between joint forces and joint accelerations. The forces acting on a joint are the sequenced joint force and linear and angular inertia of the joint.

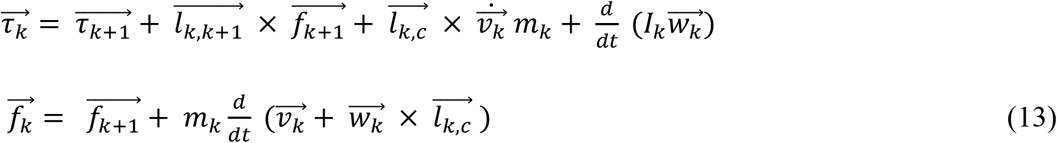

The compact form of Eq.13 can be given as in Eq.14.

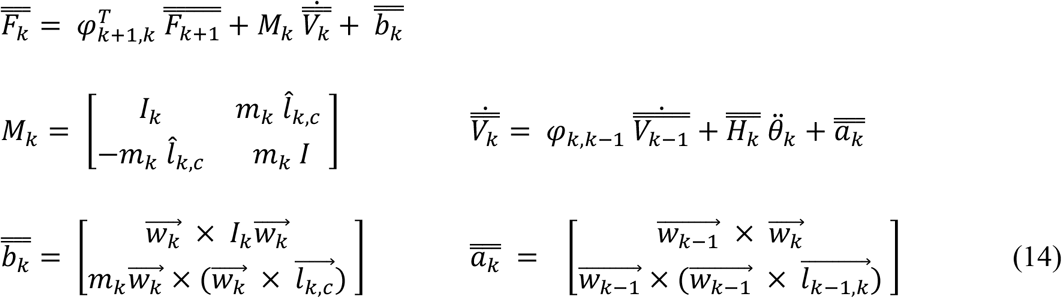

If the robot joint dynamic equations are put together, Eq.15 is obtained.

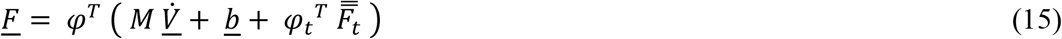

The above equation includes all forces acting on the robot joints. To reach forces that make the robot joint accelerate, the force vector *F* is multiplicated with the rotation axis matrix as in Eq.16.

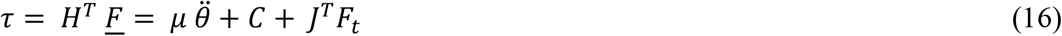

Based on MATLAB kinematic code for moving base-robot systems, a dynamic model is built. The torque expressions are related to Jacobian terms. Because of that dynamic model code can be easily improved by starting with kinematic code. The first addition to kinematic code is creating the *μ* matrix according to base velocities, base accelerations, and fixed base dual robot “*μ*”. The fixed base dual robot “*μ*” requires robot joint masses, inertia tensors, position vector of the center of mass, and joint angular velocity vectors. Masses, inertia tensors, and center of gravities concerning joint frames are found by investigating the Solidworks drawing of the robot arm. Joint angular velocity vectors are provided from the previous iteration with zero initial points.

After evaluating the fixed base “*μ*”, the moving base features are added to the “*μ*” and a new “*μ*” is created. These additions are base mass matrix which depends on base mass, inertia tensor, the center of gravity, and base-robot mass matrices. C matrix is created based on H, *φ*, a vector, b vector, and mass matrix. After calculating these matrices, the forward dynamics of the robot with a moving base are simulated. Above torque, expression is prepared as torque input and acceleration output given in Fig.5 and Eq.17.

**Figure 5:**
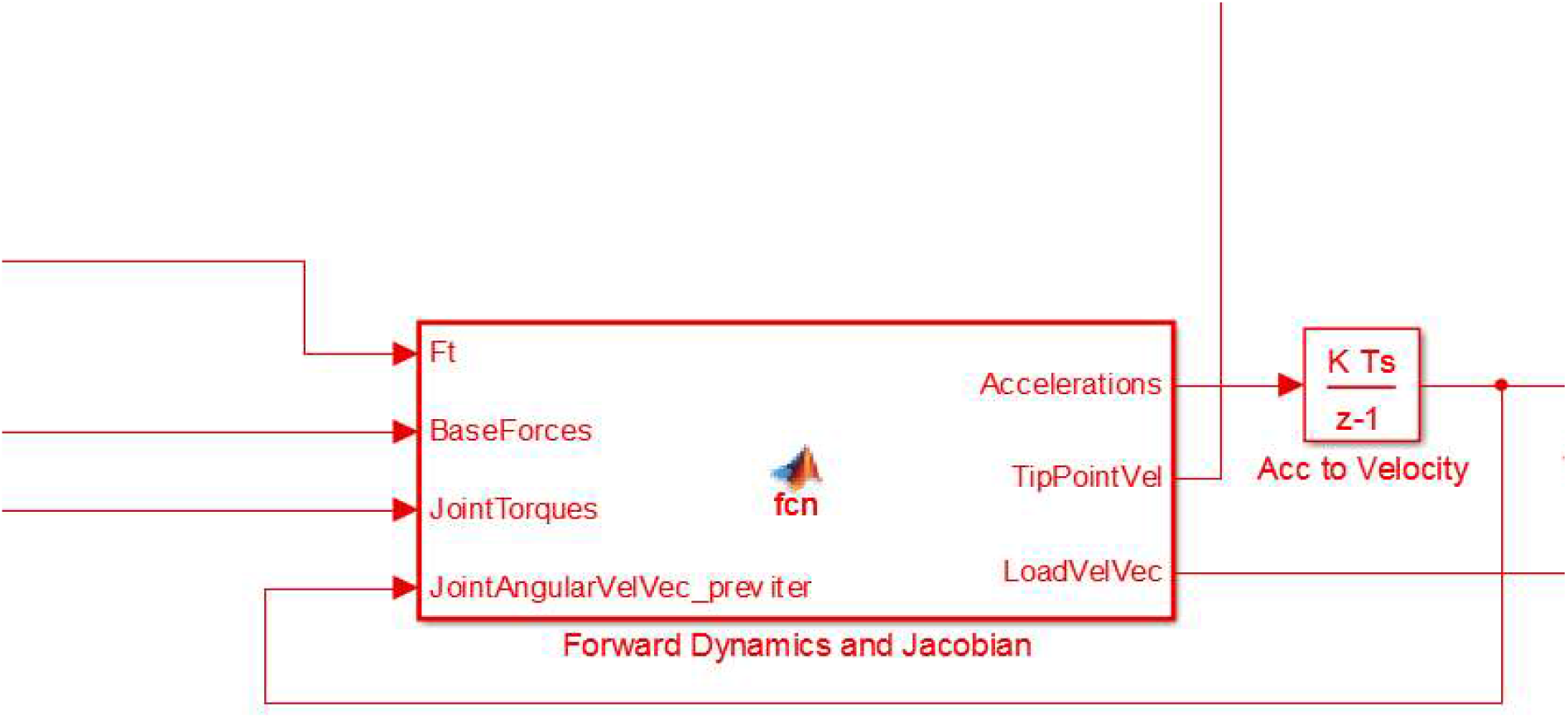
Simulink diagram of the forward dynamics.

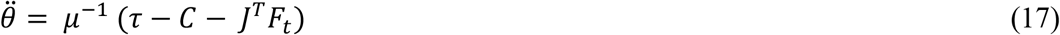

The inputs are; Tip Forces (12×1), Base Forces(6×1), Joint Torques (12×1), and Joint Angular Velocity Vectors (12×1), respectively. The outputs are; Accelerations (18×1), Tip Point Velocities (12×1), and Load Velocity (6×1), respectively.

## 3. Results

After applying the adaptive AR-based PDC feature extraction method to the train data matrix for each class, the time-variant estimated MVAR parameters were obtained. The size of the parameter vector for each class is 32000 × 200. The rows represent time in terms of data samples, and the columns show the parameter values estimated over samples. The most significant values of the adaptive PDC measures at a 99% level of significance after applying the surrogate data method are given in Fig.6 for each class of wrist movement states.

**Figure 6:**
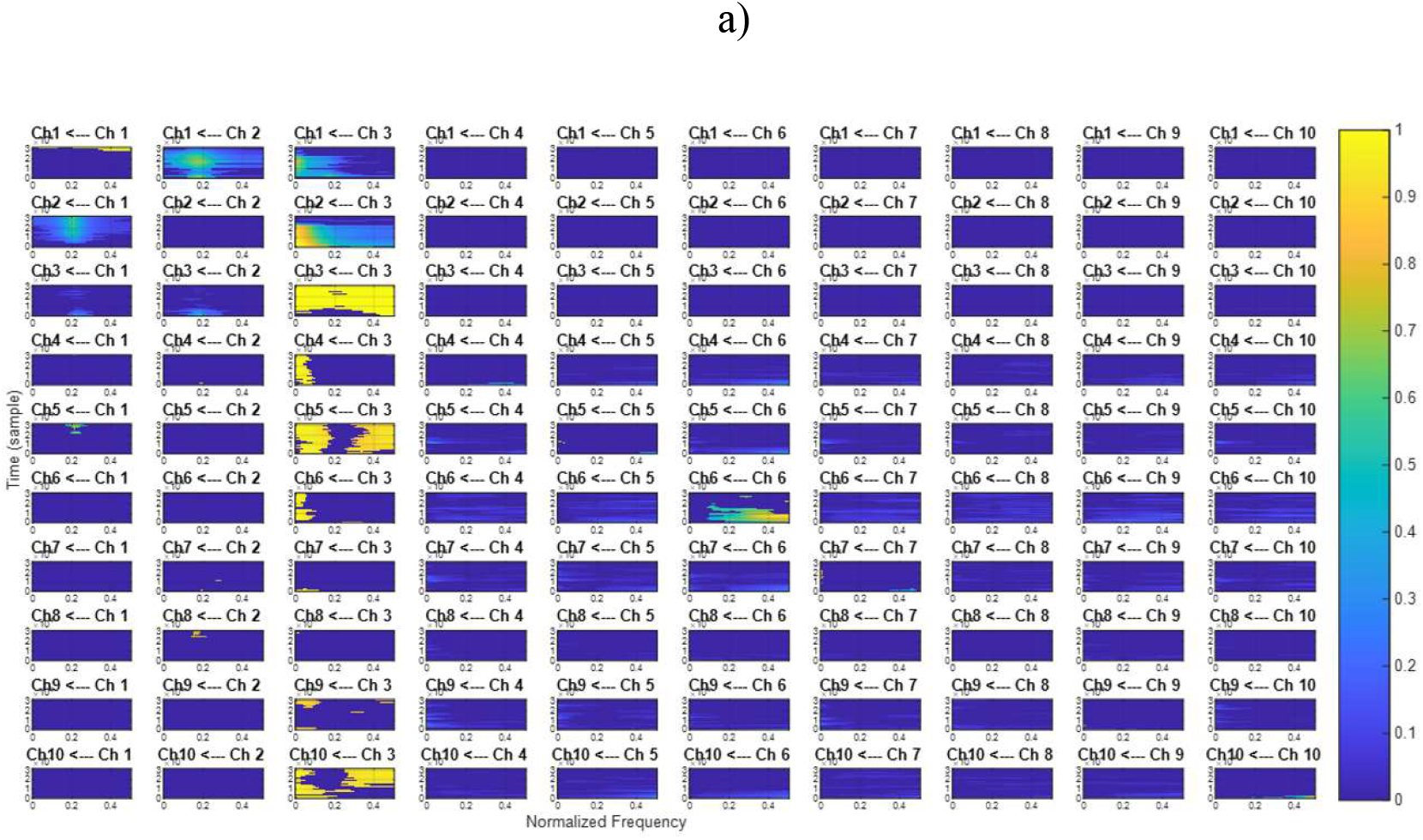
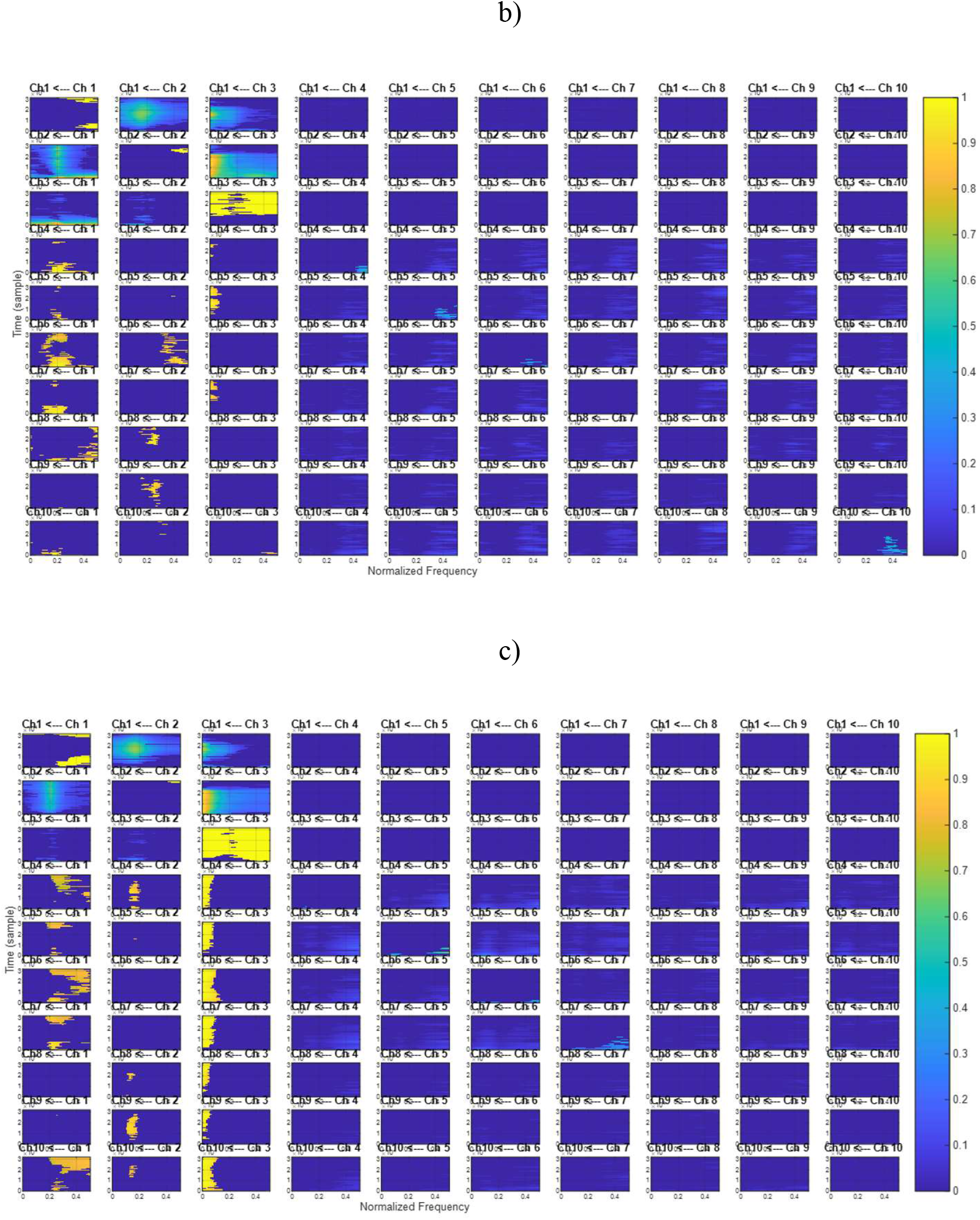
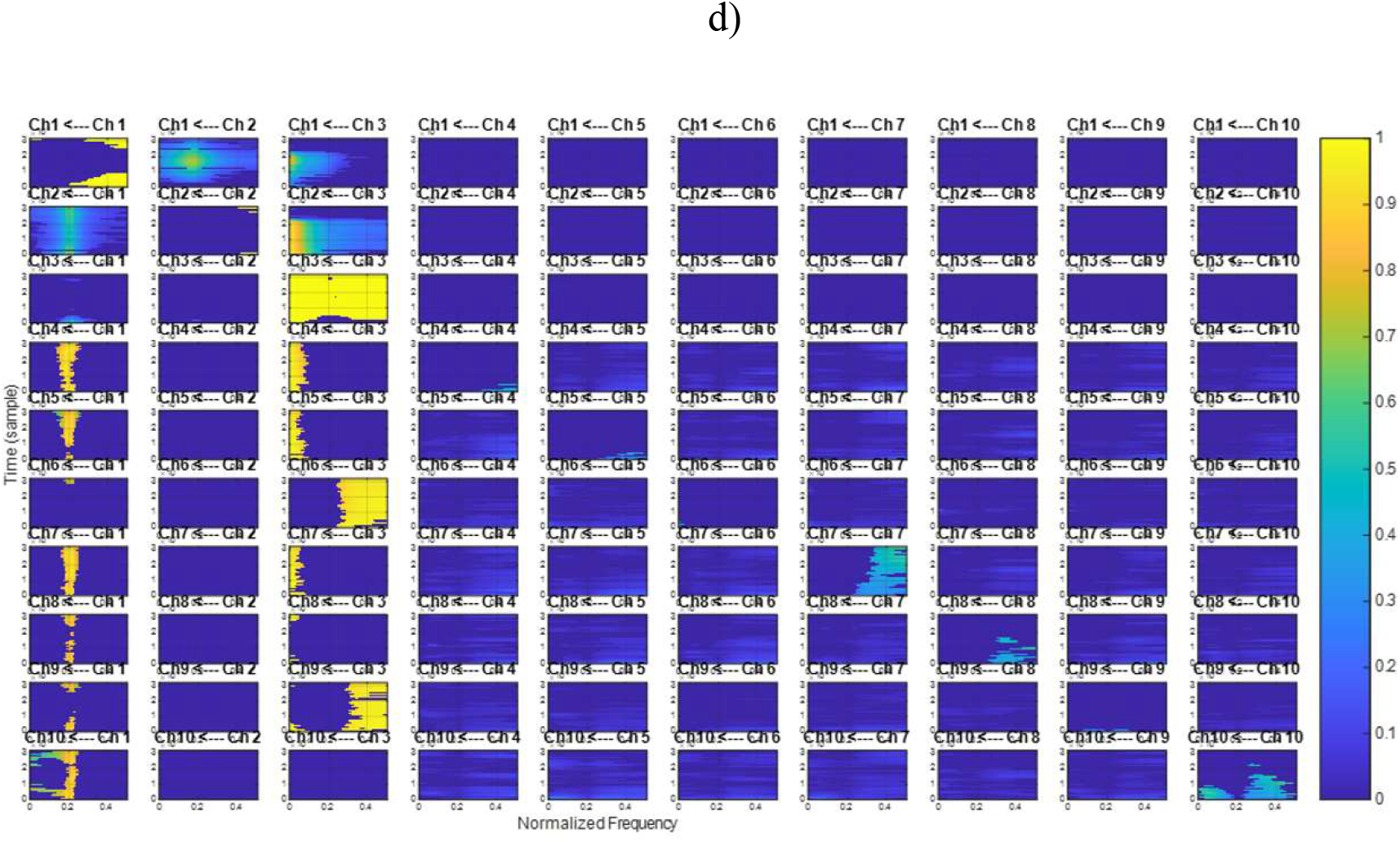
PDC for the simulated model utilizing the Kalman filtering approach. The x-axis represents normalized frequency ([0 0.5] corresponding to [0 *F*_s_/2]) and the y-axis represents the time direction in terms of data samples. Wrist movement states: a) right, b) forward, c) left, and d) backward.

The adaptive PDC results show the connectivity between channels. It can be inferred from Fig.6, that the most significant direct coupling activities emerge from channel 3 to channels 1, 2, 4, and 5, respectively. The model parameters are accurately tracked and each class has different patterns.

The adaptive AR-based PDC feature matrices are concatenated and the feature train matrix is built, the obtained train matrix is fed into the ELM. The size of the feature train matrix is 128000 × 200.

A principle component analysis-based dimension reduction algorithm, which is an orthogonal transformation constructing the relevant features, is applied to the feature train matrix because the number of extracted features is too much. The first four components having the most variance in the feature data are selected. After the dimensionality reduction process, the size of the feature train matrix becomes 128000 × 4.

The obtained data are randomly divided into training, testing, and validation sets. Every time the system is executed samples were used for each task (70400 samples (55%) are used for training, 25600 samples are used for validation (20%) and the remaining 32000 trials were used for the test (25%)). ELM has trained for 100 epochs (iterations) by incrementing the number of neurons in the hidden layer (hidden node). For each hidden node, the classification accuracy was evaluated in terms of the “root mean squared error” criteria. Fig.7 represents the performance curve sensitivity concerning the hidden node.

**Figure 7:**
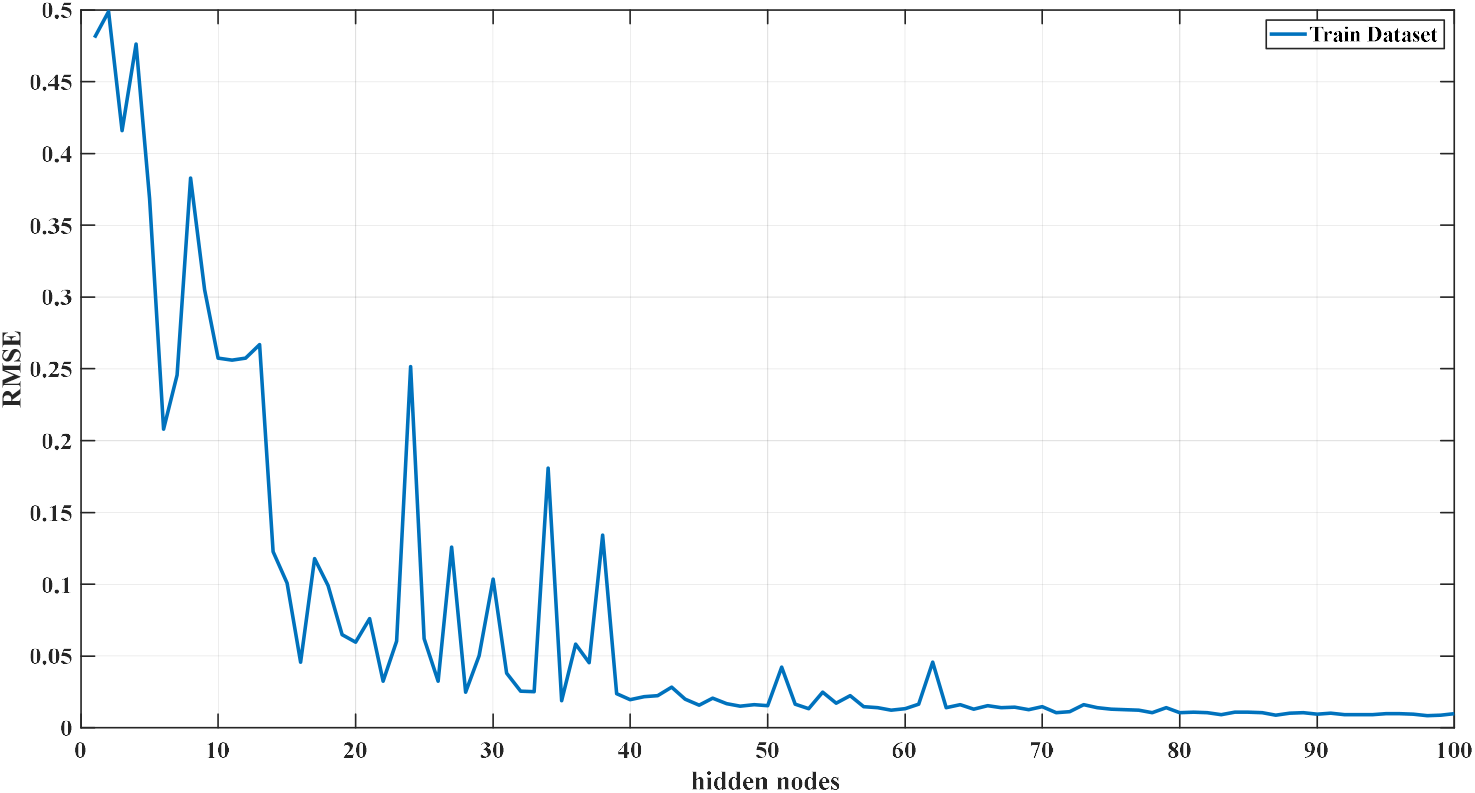
Hidden node selecting criteria.

The rmse error approaches the equilibrium state while increasing the hidden nodes. Therefore, the number of hidden neurons is chosen as 100. The training ratio is selected using a trial & error process as 0.7. The activation function of the output layer is chosen as “sigmoid”.

The confusion matrices are given in Fig.8.

**Figure 8:**
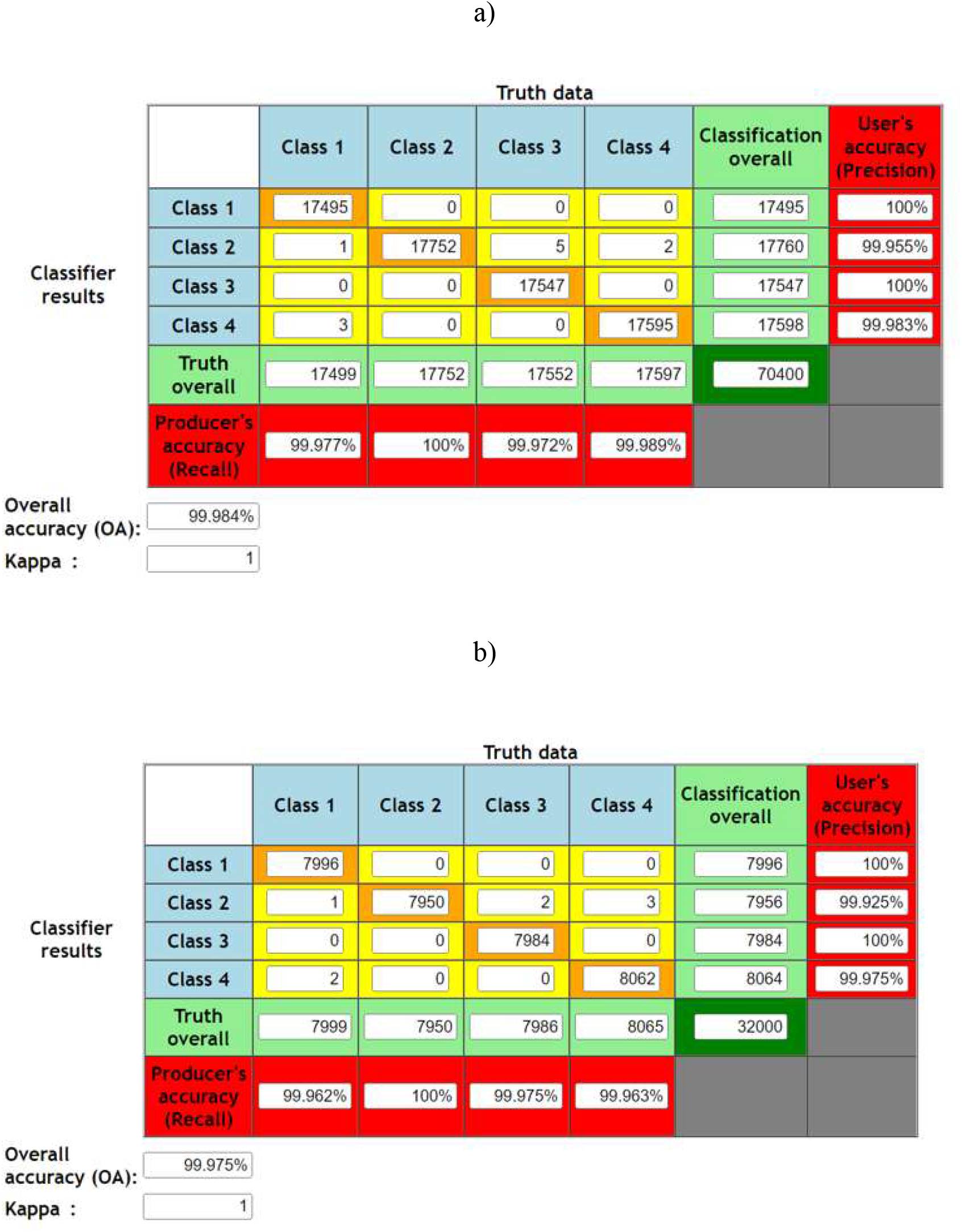
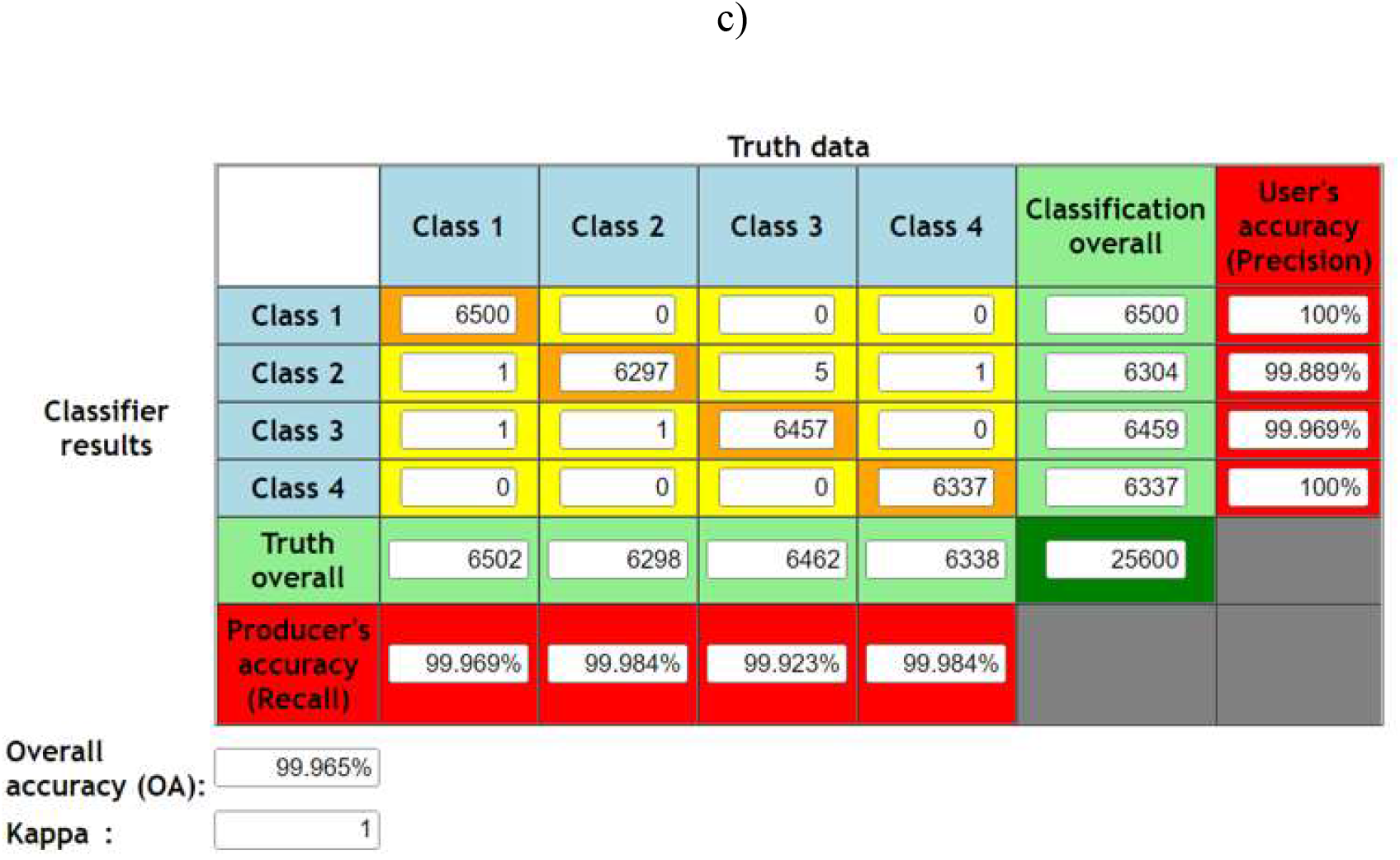
Confusion matrices a) training b) test c) validation. Class 1: right, Class 2: forward, Class 3: left, Class 4: backward

Test data is used for the external validation process. The size of the reduced feature test matrix is 58400 × 4. After feeding the test data to the trained ELM model, the external validation accuracy of the ELM is found as 84.88%. A comparative analysis was conducted over the external validation results using various classification methods and given in Table 1.

**Table 1:** Comparison of external validation classification results. (Abbreviations; SVM: Support Vector Machine, ELM: Extreme Learning Machine, NB: Naive Bayesian, k-NN: k-Nearest Neighbor, ANN: Artificial Neural Network).

| **Classifier** | **Sensitivity [%]** | **Precision [%]** | **Specificity [%]** | **F-measure [%]** | **Accuracy[%]** |
| --- | --- | --- | --- | --- | --- |
| **SVM** | 83.6 | 82.88 | 82.71 | 83.25 | 83.17 |
| **ELM** | 85.12 | 83.73 | 83.62 | 84.95 | 84.88 |
| **NB** | 75.45 | 77.03 | 76.74 | 76.23 | 76.08 |
| **k-NN** | 78.54 | 67.93 | 76.79 | 78.23 | 77.71 |
| **ANN** | 79.78 | 79.23 | 76.79 | 79.50 | 78.43 |

According to the external validation results, the ELM outperforms the other machine learning models, and also the computational speed is extremely fast.

## 4. Discussion and Conclusive Summary

The obtained results clarify that by applying the proposed methodology, control signals from an individual’s MEG signals can control the robotic arm. Time-varying cortical neural connectivity features are strong biomarkers for characterizing the MEG patterns, which originated from wrist movements. According to the external validation classification results, ELM outperforms the other classifiers by achieving the highest accuracy. The accuracy of which these control signals are extracted from the MEG signals is extensively measured using the wide industry-employed receiver operator characteristic curves. After the signal processing part is taken care of, it is necessary to establish the dynamic model of the robot, create and simulate the solid model, eliminate the errors caused by the system dynamics in the robot joints, and synchronize with the brain signals. MEG-based BCI systems are capable to be robustly utilized as a control device through the proposed framework.

## Patient-informed consent

There is no need for patient-informed consent.

## Ethics committee approval

There is no need for ethics committee approval.

## Author contribution subject and rate

**CU (100%):** Design the research, data collection, and analyses and wrote the whole manuscript.

**CU (100%):** Organized the research and supervised the article write-up.

**CU (100%):** Contributed with comments on research design and slide interpretation.

**CU (100%):** Contributed with comments on manuscript organization and write-up.

## References

[1] Mellinger, J., Schalk, G., Braun, C., Preissl, H., Rosenstiel, W., Birbaumer, N., & Kübler, A. (2007). An Meg-based brain–computer interface (BCI). NeuroImage, 36(3), 581–593. https://doi.org/10.1016/j.neuroimage.2007.03.019

[2] Gutiérrez David, Corral, G. H., & Zentina, L. M. (2008). Using EEG/MEG data of cognitive processes in brain-computer interfaces. AIP Conference Proceedings. https://doi.org/10.1063/1.2979300

[3] Çağlayan, O., & Arslan, R. B. (2013). Robotic arm control with brain computer interface using P300 and SSVEP. Biomedical Engineering. https://doi.org/10.2316/p.2013.791-082

[4] Nandikolla, V., & Medina Portilla, D. A. (2022). Teleoperation robot control of a hybrid EEG-based BCI arm manipulator using Ros. Journal of Robotics, 2022, 1–14. https://doi.org/10.1155/2022/5335523

[5] Qasim, M., & Ismael, O. Y. (2022). Shared control of a robot arm using BCI and Computer Vision. Journal Européen Des Systèmes Automatisés, 55(1), 139–146. https://doi.org/10.18280/jesa.550115

[6] Musk, E. (2019). An integrated brain-machine interface platform with thousands of channels. Journal of Medical Internet Research, 21(10). https://doi.org/10.2196/16194

[7] Spüler, M., Rosenstiel, W., & Bogdan, M. (2012). Adaptive SVM-based classification increases performance of a Meg-based brain-computer interface (BCI). Artificial Neural Networks and Machine Learning – ICANN 2012, 669–676. https://doi.org/10.1007/978-3-642-33269-2_84

[8] Rezaei, S., Tavakolian, K., Nasrabadi, A. M., & Setarehdan, S. K. (2006). Different classification techniques considering brain computer interface applications. Journal of Neural Engineering, 3(2), 139–144. https://doi.org/10.1088/1741-2560/3/2/008

[9] Corsi, M.-C., Chavez, M., Schwartz, D., Hugueville, L., Khambhati, A. N., Bassett, D. S., & De Vico Fallani, F. (2019). Integrating EEG and MEG signals to improve motor imagery classification in brain–computer interface. International Journal of Neural Systems, 29(01), 1850014. https://doi.org/10.1142/s0129065718500144

[10] Sabra, N. I., & Abdel Wahed, M. (2011). The use of Meg-based brain computer interface for classification of wrist movements in four different directions. 2011 28th National Radio Science Conference (NRSC). https://doi.org/10.1109/nrsc.2011.5873644

[11] Daliri, M. R. (2014). A hybrid method for the decoding of spatial attention using the Meg Brain Signals. Biomedical Signal Processing and Control, 10, 308–312. https://doi.org/10.1016/j.bspc.2012.12.005

[12] Scherer, R., & Vidaurre, C. (2018). Motor imagery based brain–computer interfaces. Smart Wheelchairs and Brain-Computer Interfaces, 171–195. https://doi.org/10.1016/b978-0-12-812892-3.00008-x

[13] Uyulan, C., & Erguzel, T. T. (2017). Analysis of time – frequency EEG feature extraction methods for Mental Task Classification. International Journal of Computational Intelligence Systems, 10(1), 1280. https://doi.org/10.2991/ijcis.10.1.87

[14] Uyulan, C., & Erguzel, T. (2016). Comparison of wavelet families for Mental Task Classification. The Journal of Neurobehavioral Sciences, 3(2), 59. https://doi.org/10.5455/jnbs.1454666348

[15] Sun, L., & Feng, Z. R. (2016). Classification of imagery motor EEG data with wavelet denoising and features selection. 2016 International Conference on Wavelet Analysis and Pattern Recognition (ICWAPR). https://doi.org/10.1109/icwapr.2016.7731641

[16] Murugappan, M., & Murugappan, S. (2013). Human emotion recognition through short time electroencephalogram (EEG) signals using Fast Fourier transform (FFT). 2013 IEEE 9th International Colloquium on Signal Processing and Its Applications. https://doi.org/10.1109/cspa.2013.6530058

[17] Zaveri, H. P., Duckrow, R. B., & Spencer, S. S. (2006). On the use of bipolar montages for time-series analysis of intracranial electroencephalograms. Clinical Neurophysiology, 117(9), 2102–2108. https://doi.org/10.1016/j.clinph.2006.05.032

[18] Ludwig, K. A., Miriani, R. M., Langhals, N. B., Joseph, M. D., Anderson, D. J., & Kipke, D. R. (2009). Using a common average reference to improve cortical neuron recordings from microelectrode arrays. Journal of Neurophysiology, 101(3), 1679–1689. https://doi.org/10.1152/jn.90989.2008

[19] Mourino, J., del R Millan, J., Cincotti, F., Chiappa, S., Jane, R., & Babiloni, F. (2001). Spatial filtering in the training process of a brain computer interface. 2001 Conference Proceedings of the 23rd Annual International Conference of the IEEE Engineering in Medicine and Biology Society. https://doi.org/10.1109/iembs.2001.1019016

[20] Saha, P. K., Rahman, M. A., Alam, M. K., Ferdowsi, A., & Mollah, M. N. (2021). Common spatial pattern in frequency domain for feature extraction and classification of multichannel EEG signals. SN Computer Science, 2(3). https://doi.org/10.1007/s42979-021-00586-9

[21] Uyulan, C., Ergüzel, T. T., & Tarhan, N. (2019). Entropy-based feature extraction technique in conjunction with Wavelet packet transform for multi-mental task classification. Biomedical Engineering / Biomedizinische Technik, 64(5), 529–542. https://doi.org/10.1515/bmt-2018-0105

[22] Güçlü, U., Güçlütürk, Y., & Loo, C. K. (2011). Evaluation of fractal dimension estimation methods for feature extraction in motor imagery based brain computer interface. Procedia Computer Science, 3, 589–594. https://doi.org/10.1016/j.procs.2010.12.098

[23] Aznan, Nik Khadijah Nik & Yang, Yeon-Mo. (2013). Applying Kalman filter in EEG-based brain computer interface for motor imagery classification. 2013 International Conference on ICT Convergence (ICTC). https://doi.org/10.1109/ictc.2013.6675451

[24] Nicolas-Alonso, L. F., & Gomez-Gil, J. (2012). Brain Computer Interfaces, a review. Sensors, 12(2), 1211–1279. https://doi.org/10.3390/s120201211

[25] Madi, M. K., & Karameh, F. N. (2017). Hybrid cubature kalman filtering for identifying nonlinear models from sampled recording: Estimation of neuronal dynamics. PLOS ONE, 12(7). https://doi.org/10.1371/journal.pone.0181513

[26] Xin Ma. (2011). The research of brain-computer interface based on AAR parameters and neural networks classifier. Proceedings of 2011 International Conference on Computer Science and Network Technology. https://doi.org/10.1109/iccsnt.2011.6182491

[27] Nilesh, R., & Sunil, W. (2021). Improving extreme learning machine through optimization a review. 2021 7th International Conference on Advanced Computing and Communication Systems (ICACCS). https://doi.org/10.1109/icaccs51430.2021.9442007

[28] Lefebvre, G., & Cumin, J. (2016). Recognizing human actions based on Extreme Learning Machines. Proceedings of the 11th Joint Conference on Computer Vision, Imaging and Computer Graphics Theory and Applications. https://doi.org/10.5220/0005675004780483

[29] Mukherjee, H., Das, S., Ghosh, S., Obaidullah, S. M., Santosh, K. C., Das, N., & Roy, K. (2019). A study on the Extreme Learning Machine and its applications. Document Processing Using Machine Learning, 43–52. https://doi.org/10.1201/9780429277573-4

[30] Matias, T., Souza, F., Araújo, R., Gonçalves, N., & Barreto, J. P. (2015). On-line sequential extreme learning machine based on recursive partial least squares. Journal of Process Control, 27, 15–21. https://doi.org/10.1016/j.jprocont.2015.01.004

[31] Huang, G.-B., Chen, L., & Siew, C.-K. (2006). Universal approximation using incremental constructive feedforward networks with random hidden nodes. IEEE Transactions on Neural Networks, 17(4), 879–892. https://pubmed.ncbi.nlm.nih.gov/16856652/

[32] Rong, H.-J., Ong, Y.-S., Tan, A.-H., & Zhu, Z. (2008). A fast pruned-extreme learning machine for classification problem. Neurocomputing, 72(1-3), 359–366. https://doi.org/10.1016/j.neucom.2008.01.005

[33] Feng, G. Guang-Bin Huang, Qingping Lin, & Gay, R. (2009). Error minimized extreme learning machine with growth of hidden nodes and incremental learning. IEEE Transactions on Neural Networks, 20(8), 1352–1357. https://doi.org/10.1109/tnn.2009.2024147

[34] Silva, D. N., Pacifico, L. D., & Ludermir, T. B. (2011). An evolutionary extreme learning machine based on group search optimization. 2011 IEEE Congress of Evolutionary Computation (CEC). https://doi.org/10.1109/cec.2011.5949670

[35] Lan, Y., Soh, Y. C., & Huang, G.-B. (2010). Two-stage extreme learning machine for regression. Neurocomputing, 73(16-18), 3028–3038. https://doi.org/10.1016/j.neucom.2010.07.012

[36] Deng, W., Zheng, Q., & Chen, L. (2009). Regularized extreme learning machine. 2009 IEEE Symposium on Computational Intelligence and Data Mining. https://doi.org/10.1109/cidm.2009.4938676

[37] Ding, X.-jian, Liu, X.-guang, & Xu, X. (2016). An optimization method of extreme learning machine for regression. Proceedings of the 31st Annual ACM Symposium on Applied Computing. https://doi.org/10.1145/2851613.2851882

[38] Ding, S., Zhang, N., Xu, X., Guo, L., & Zhang, J. (2015). Deep Extreme Learning Machine and its application in EEG classification. Mathematical Problems in Engineering, 2015, 1–11. https://doi.org/10.1155/2015/129021

[39] Stosic, D., Stosic, D., & Ludermir, T. (2016). Voting based Q-Generalized Extreme Learning Machine. Neurocomputing, 174, 1021–1030. https://doi.org/10.1016/j.neucom.2015.10.028

[40] Omidvarnia, A., Mesbah, M., O’Toole, J. M., Colditz, P., & Boashash, B. (2011). Analysis of the time-varying cortical neural connectivity in the newborn EEG: A Time-frequency approach. International Workshop on Systems, Signal Processing and Their Applications, WOSSPA. https://doi.org/10.1109/wosspa.2011.5931445

[41] Winterhalder, M., Schelter, B., Hesse, W., Schwab, K., Leistritz, L., Klan, D., Bauer, R., Timmer, J., & Witte, H. (2005). Comparison of linear signal processing techniques to infer directed interactions in multivariate neural systems. Signal Processing, 85(11), 2137–2160. https://doi.org/10.1016/j.sigpro.2005.07.011

[42] Arnold, M., Milner, X. H. R., Witte, H., Bauer, R., & Braun, C. (1998). Adaptive AR modeling of nonstationary time series by means of Kalman filtering. IEEE Transactions on Biomedical Engineering, 45(5), 553–562. https://doi.org/10.1109/10.668741

[43] Fan Wang, & Balakrishnan, V. (2002). Robust kalman filters for linear time-varying systems with stochastic parametric uncertainties. IEEE Transactions on Signal Processing, 50(4), 803–813. https://doi.org/10.1109/78.992124

[44] Pagnotta, M. F., Plomp, G., & Pascucci, D. (2019). A regularized and smoothed general linear Kalman filter for more accurate estimation of time-varying directed connectivity. 2019 41st Annual International Conference of the IEEE Engineering in Medicine and Biology Society (EMBC). https://doi.org/10.1109/embc.2019.8857915

[45] Pascucci, D., Rubega, M., & Plomp, G. (2020). Modeling time-varying brain networks with a self-tuning optimized Kalman filter. PLOS Computational Biology, 16(8). https://doi.org/10.1371/journal.pcbi.1007566

[46] Ghumare, E. G., Schrooten, M., Vandenberghe, R., & Dupont, P. (2018). A time-varying connectivity analysis from distributed EEG SOURCES: A simulation study. Brain Topography, 31(5), 721–737. https://doi.org/10.1007/s10548-018-0621-3

[47] Uyulan, C., de la Salle, S., Erguzel, T. T., Lynn, E., Blier, P., Knott, V., Adamson, M. M., Zelka, M., & Tarhan, N. (2021). Depression diagnosis modeling with advanced computational methods: Frequency-domain emvar and deep learning. Clinical EEG and Neuroscience, 53(1), 24–36. https://doi.org/10.1177/15500594211018545

[48] Hu, J., Zhang, J., Zhang, C., & Wang, J. (2016). A new deep neural network based on a stack of single-hidden-layer feedforward neural networks with randomly fixed hidden neurons. Neurocomputing, 171, 63–72. https://doi.org/10.1016/j.neucom.2015.06.017

[49] Paul, I., Ghosh, S., & Konar, A. (2020). Voice Command decoding for position control of Jaco robot arm using a type-2 fuzzy classifier. 2020 IEEE International Conference on Electronics, Computing and Communication Technologies (CONECCT). https://doi.org/10.1109/conecct50063.2020.9198684

